# Natural variation in *Caenorhabditis elegans* responses to the anthelmintic emodepside

**DOI:** 10.1101/2021.01.05.425329

**Authors:** Janneke Wit, Briana C. Rodriguez, Erik. C. Andersen

**Author notes:** **Corresponding Author:** Erik C. Andersen, Ph.D., Department of Molecular Biosciences, Northwestern University, 4619 Silverman Hall, 2205 Tech Drive, Evanston, IL 60208. **Journal:** *IJP DDR^1^.

## Abstract

Treatment of parasitic nematode infections depends primarily on the use of anthelmintics. However, this drug arsenal is limited, and resistance against most anthelmintics is widespread. Emodepside is a new anthelmintic drug effective against gastrointestinal and filarial nematodes. Nematodes that are resistant to other anthelmintic drug classes are susceptible to emodepside, indicating that the emodepside mode of action is distinct from previous anthelmintics. The laboratory-adapted *Caenorhabditis elegans* strain N2 is sensitive to emodepside, and genetic selection and *in vitro* experiments implicated *slo-1*, a BK potassium channel gene, in emodepside mode of action. In an effort to understand how natural populations will respond to emodepside, we measured brood sizes and developmental rates of wild *C. elegans* strains after exposure to the drug and found natural variation across the species. Some of the observed variation in *C. elegans* emodepside responses correlates with amino acid substitutions in *slo-1*, but genetic mechanisms other than *slo-1* coding variants likely underlie emodepside resistance in wild *C. elegans* strains. Additionally, the assayed strains have higher offspring production in low concentrations of emodepside (a hormetic effect), which could impact treatment strategies when parasites are underdosed. We find that natural variation affects emodepside sensitivity, supporting the suitability of *C. elegans* as a model system to study emodepside responses across natural nematode populations.

**Graphical abstract:** 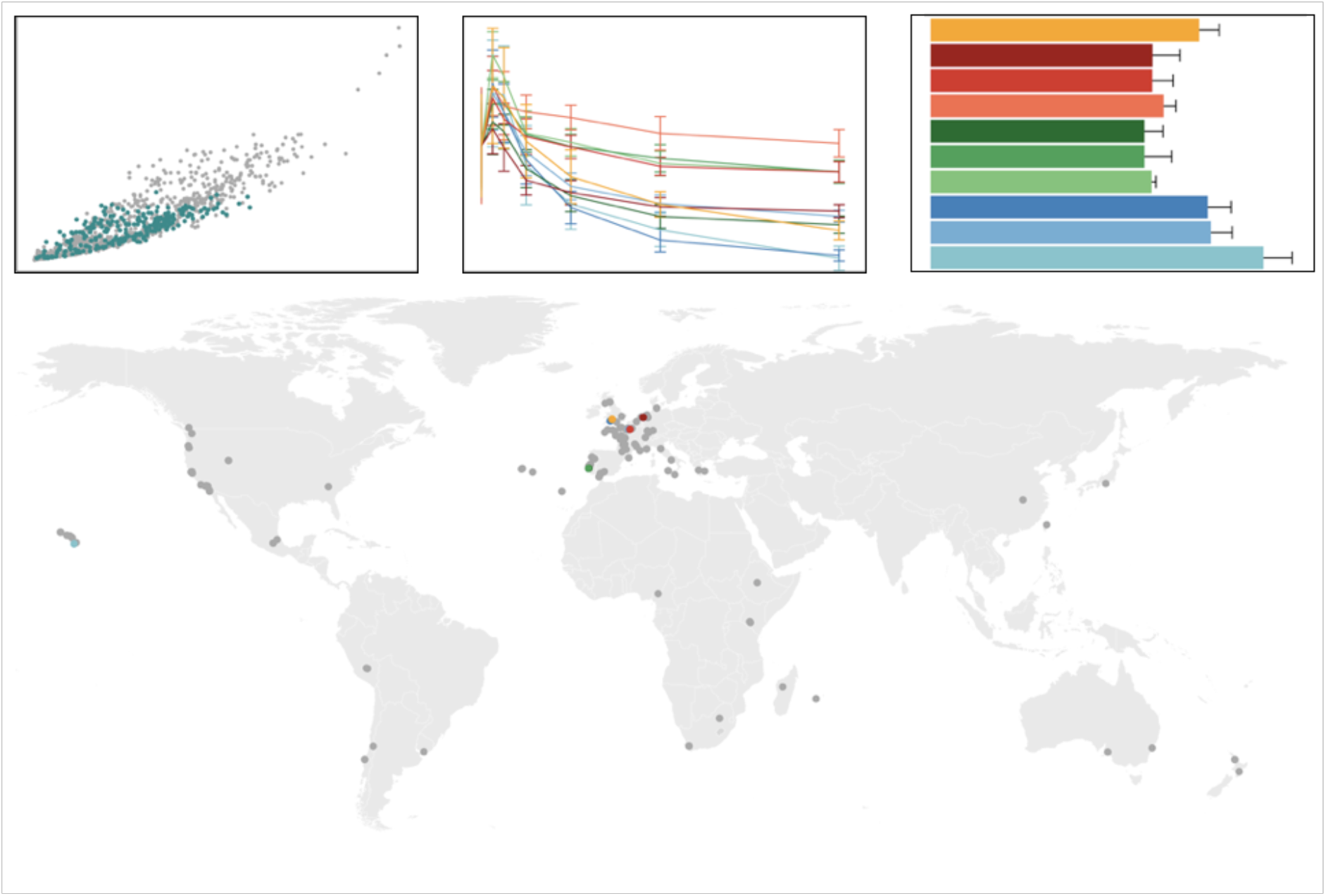

**Highlights:**

- Emodepside responses vary across the *C. elegans* species.
- Wild strains of *C. elegans* model natural differences in parasite emodepside responses.
- Variation in the emodepside target *slo-1* and other loci correlate with resistance.
- Low doses of emodepside cause a hormetic effect on offspring production.

## 1. Introduction

Helminth infections are a major threat to animal and human health, and control measures depend heavily on a small arsenal of anthelmintic drugs. Resistance against most anthelmintic drug classes is widespread and documented for several species (Kotze and Prichard, 2016; McKellar and Jackson, 2004). New anthelmintics with a distinct mode of action can be used to treat populations resistant to multiple anthelmintics, but the introduction of new compounds is rare (Epe and Kaminsky, 2013). One of the newest anthelmintics, the cyclooctadepsipeptide (COPD) emodepside, has been commercially available since 2007 (Epe and Kaminsky, 2013). It is a semisynthetic derivative of a natural metabolite from the fungus *Mycelia sterilia* (Harder and von Samson-Himmelstjerna, 2001). As a broad spectrum anthelmintic, emodepside is efficacious against gastrointestinal nematodes and filarial nematodes (Harder et al., 2003; Zahner et al., 2001) and is currently approved for treatment of helminth infections of cats and dogs in combination with praziquantel (Altreuther et al., 2005). Field resistance has not been reported since its introduction (Prichard, 2017). Importantly, emodepside is effective against multi-drug resistant parasitic nematode strains, including ivermectin- and levamisole-resistant *Haemonchus contortus* (Harder et al., 2005; von Samson-Himmelstjerna et al., 2005).

Responses to COPD have been studied in both parasitic nematodes and the free-living nematode *Caenorhabditis elegans*. Initial *in vitro* studies using *Ascaris suum* suggested that the COPD PF1022A, the parent compound in emodepside synthesis (Jeschke et al., 2005), displaces GABAergic ligands from somatic muscle preparations (Harder et al., 2005). However, later work comparing the effect of GABA and emodepside on the rate of relaxation of contracted *A. suum* muscle showed that emodepside does not act directly on the GABAergic pathway (Willson et al., 2003; Willson J, Holden-Dye L, Harder A, Walker RJ, 2001). Another promising lead was the identification of a putative target protein, HC110-R, from a *H. contortus* cDNA library (Saeger et al., 2001). Alignment revealed HC110-R had 48% identity and 76% similarity to the *C. elegans* latrophilin receptor LAT-1. Although predicted to be a heptahelical transmembrane protein, the exact function of HC110-R is unknown (Mühlfeld et al., 2009). Latrophilin is a G protein-coupled receptor in the secretin receptor family and a Ca^2+^-independent receptor of alpha-latrotoxin (WeIz et al., 2005). *C. elegans* larvae express *lat-1* in pharyngeal muscle, and adults express it in both pharyngeal and non-pharyngeal neurons (Willson et al., 2004). In the laboratory strain N2, emodepside inhibits pharyngeal pumping, egg-laying, as well as locomotion (Bull et al., 2007). Putative null mutations in *lat-1* are less sensitive to emodepside-induced inhibition of pharyngeal pumping, but locomotor activity is inhibited (Guest et al., 2007; Willson et al., 2004). This inhibition of locomotion suggests that emodepside affects additional pathways independent of *lat-1*.

A subsequent mutagenesis screen using *C. elegans* identified mutations in the Ca^2+^-activated K^+^ channel (BK-channel) gene *slo-1* in nine emodepside resistant mutants (Guest et al., 2007). These mutants were identified as highly resistant to inhibition of both pharyngeal pumping and locomotor activity by emodepside. Gain-of-function mutations in *slo-1* show decreased locomotion and pharyngeal pumping similar to emodepside-treated nematodes (Davies et al., 2003), suggesting that emodepside activates SLO-1 signaling. Additionally, a putative *slo-1* null allele, *slo-1(js379)*, responded to emodepside treatment like mutants from the screen (Guest et al., 2007). Tissue-specific rescue experiments in the putative *slo-1* null background showed that emodepside inhibited locomotion by *slo-1* expressed in both neurons and body wall muscle (Guest et al., 2007). However, feeding was inhibited by emodepside effects on pharyngeal-specific neurons alone and not through muscle. Subsequently, emodepside was shown to open SLO-1 channels expressed in *Xenopus laevis* oocytes (Kulke et al., 2014). Taken together, these results suggest that emodepside acts mainly through a *slo-1* dependent pathway, and that the drug opens SLO-1 channels to inhibit locomotion and pharyngeal pumping in *C. elegans*.

The above studies on emodepside mode of action and resistance in *C. elegans* focused on the N2 laboratory strain and mutants in that genetic background. Although *C. elegans* is a great model organism for parasitic nematodes (Bürglin et al., 1998; Dilks et al., 2020; Hahnel et al., 2018), studies that use only a single strain can be biased by rare variation or genetic modifiers specific to a single genetic background (Sterken et al., 2015). The observation that emodepside affects multiple nematode species suggests that its mode of action is conserved throughout the phylum. It is unlikely that one *C. elegans* strain represents all possible genes and variants that contribute to emodepside sensitivity. The use of multiple isolates in drug response studies increases the likelihood of elucidating mechanisms of resistance and drug mode of action shared by multiple strains and species (Hahnel et al., 2020; Wit et al., 2020). Natural variation across the *C. elegans* species is archived in the *C. elegans* Natural Diversity Resource (CeNDR) (Cook et al., 2017) and offers a powerful approach to look for genetic variation that underlies the different responses to emodepside, as has been done for other drugs (Brady et al., 2019; Evans and Andersen, 2020; Hahnel et al., 2018; Zamanian et al., 2018; Zdraljevic et al., 2019, 2017).

Here, we measured emodepside responses in a set of *C. elegan*s wild strains to demonstrate that the effect of this anthelmintic on development and brood size depends on the genetic background. Across a set of nine wild strains and the laboratory strain N2, we show that natural coding variation in *slo-1* is correlated with differences in response to emodepside, but that additional variation impacts emodepside responses. This result illustrates the need for broader comparisons of anthelmintic resistance within a species, as variation in genes other than *slo-1* might affect emodepside susceptibility. Additionally, it highlights the power of using *C. elegans* natural variation for studies of emodepside mode of action and resistance because this variation might recapitulate diversity present in parasite populations.

## 2. Materials and Methods

### 2.1 Strains

Animals were maintained at 20°C on modified nematode growth medium (NGMA) containing 1% agar and 0.7% agarose seeded with the *E. coli* strain OP50 (Andersen et al., 2014). The laboratory strain N2 and a set of nine wild strains from the *C. elegans* Natural Diversity Resource (CeNDR) were used to study the response to multiple doses of emodepside and to determine the EC_50_. Additionally, two *slo-1* putative loss-of-function mutant strains, BZ142 and NM1968, were obtained from the *Caenorhabditis* Genetics Center.

### 2.2 High-throughput fitness assays

The high-throughput fitness assays (HTAs) were performed using the COPAS BIOSORT (Union Biometrica, Holliston MA) as described previously (Hahnel et al., 2018; Zdraljevic et al., 2017). In summary, the strains were grown in uncrowded conditions to avoid the induction of dauer for four generations on NMGA plates at 20°C prior to each assay. Gravid adults from the fifth generation were bleach-synchronized, and embryos were titered at one embryo per microliter of K medium (Boyd et al., 2012) into 96-well microtiter plates and incubated overnight. Hatched L1 larvae were fed with 5 mg/mL HB101 lysate (Pennsylvania State University Shared Fermentation Facility, State College, PA (García-González et al., 2017)) and cultured for 48 hours at 20°C with constant shaking. Three L4 larvae were then sorted into new microtiter plates containing K medium, 10 mg/mL HB101 lysate, 50 μM kanamycin, and either 1% DMSO or emodepside dissolved in 1% DMSO.

After sorting, animals were cultured and allowed to reproduce for 96 hours at 20°C with constant shaking. For accurate nematode length measurements, the samples were treated with sodium azide (50 mM in M9) to straighten their bodies before analysis using the COPAS BIOSORT. The COPAS BIOSORT is a large particle flow measurement device (**Figure 1**), which measures time-of-flight (TOF), extinction (EXT), and fluorescence of objects passing through the flow cell using laser beams. Animal length and optical density measure nematode development because animals get longer and more dense as they progress through development. If animals are negatively affected by emodepside, they are expected to be smaller, less optically dense, and have smaller brood sizes. Animal optical density is corrected for animal length (median.norm.EXT) for each object in each well. Object counts are used to calculate brood size (norm.n), which is the number of objects passing the laser corrected for the number of parent animals sorted into the well.

**Figure 1.**
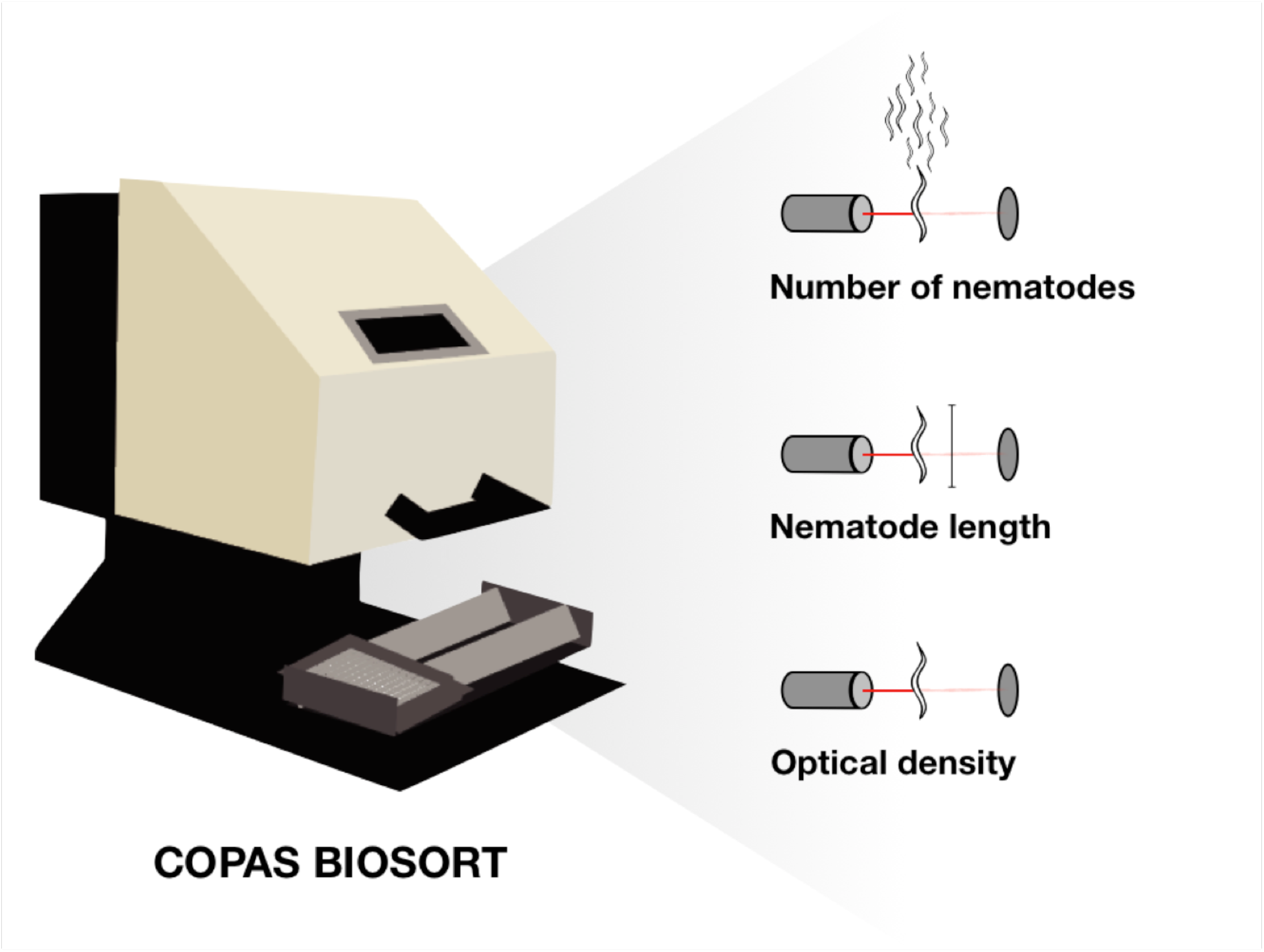
Using a COPAS BIOSORT, three independent traits were used to measure nematode responses to emodepside: brood size, nematode length (μm), and optical density.

To determine concentrations to measure differences in emodepside responses across wild strains, a dose response assay was performed using three genetically divergent *C. elegans* strains (N2, CB4856, and DL238) and four increasing concentrations of emodepside (19.6, 39.1, 78.1, and 156.3 nM). A second dose response with 9.8, 19.6, 39.1, 78.1, 156.3, and 312.5 nM emodepside was performed using nine wild strains (JU751, WN2001, NIC258, NIC265, NIC271, JU782, DL238, CB4932, and JU2586), two putative *slo-1* null mutants (BZ142 and NM1968), and the N2 strain. These 12 strains were assayed in six separate assays with four replicates in each assay. Raw phenotypic data were processed for outliers and analyzed using the R package *easysorter* (Shimko and Andersen, 2014) as described previously (Hahnel et al., 2018). For each strain, all phenotypic values were normalized by deducting the average trait value in control (DMSO) conditions.

### 2.3 Half maximal effective concentration (EC_50_) calculations

To test if emodepside had an effect on any of the three traits across the range of concentrations, extreme outliers per dose were identified and removed if values were greater or less than three times the interquartile range from the first or third quartile, respectively, using the identify_outliers from the R package *Rstatix* (Kassambara, 2020). A Kruskal-Wallis test was performed for each strain (phenotype ~ dose) using the *rstatix* package (Kassambara, 2020). For strains where emodepside had a phenotypic effect, the concentration with the highest response was determined. To calculate the half maximal effective concentration (EC_50_), the concentration at which 50% of the drug effect is reached, we could not fit from the control condition because of the hormetic effect (explained below). Instead, we fitted a linear model (developmental trait or brood size ~ dose) to the data from the dose with the peak phenotypic value to the highest concentration (312.5 nM) assayed and calculated the concentration at the midpoint of the phenotypic effect. These EC_50_ values were calculated for each strain in each of the six assays and then the mean and standard deviation of each EC_50_ were calculated.

### 2.4 Data availability

**Supplementary File 1** contains the phenotypic values for N2 and two putative *slo-1* null mutants in response to increasing concentrations of emodepside. **Supplementary File 2** contains the phenotypic values for N2, CB4856, and DL238 in response to increasing concentrations of emodepside. **Supplementary File 3** contains the raw extinction (EXT) and time of flight (TOF) measurements for N2 and two wild *C. elegans* strains (CB4856 and DL238) in control conditions and 78.1 nM emodepside. **Supplementary File 4** contains the phenotypic values for the N2 strain and nine wild *C. elegans* strains in response to increasing concentrations of emodepside. All data and scripts to generate figures can be found at https://github.com/AndersenLab/emodepside_manuscript.

## 3. Results

### 3.1 Putative *slo-1* null mutants are resistant to emodepside in the high-throughput reproduction and development assays

We assayed emodepside resistance as a function of nematode reproduction and development. These traits were measured for thousands of animals using a previously developed high-throughput assay (HTA) (see Methods, **Figure 1**) (Andersen et al., 2015; Brady et al., 2019; Evans et al., 2020, 2018; Evans and Andersen, 2020; Hahnel et al., 2018; Zamanian et al., 2018; Zdraljevic et al., 2019, 2017). In this assay, three L4 larvae were sorted into each well of a 96-well plate and allowed to grow and reproduce for 96 hours in the presence of DMSO or emodepside dissolved in DMSO. Each well contained these three parents and their offspring. After 96 hours, animal length and optical density, which are both proxies for nematode developmental stage (Andersen et al., 2015), were measured for all progeny in the well. Animals grow longer and more dense over time, and anthelmintics slow this development. Therefore, shorter and less optically dense animals after 96 hours show that emodepside had a detrimental effect on development. In addition to development, we also measured brood size as the average number of progeny produced within the 96-hour window. Although ultimate brood size and demography of the population influence statistical summaries of nematode development as measured by size (mean.TOF) or optical density (median.norm.EXT), a smaller brood size shows emodepside sensitivity (**Supplementary Figure 1**).

To confirm that our HTA could be used to quantitatively measure *C. elegans* emodepside resistance, we measured animal development and brood size for two putative *slo-1* null mutant strains, BZ142 and NM1968, and the laboratory strain, N2, across a range of concentrations. The *slo-1* mutants were shown previously to be resistant to emodepside based on locomotion and pharyngeal pumping assays (Guest et al., 2007), and the N2 strain is known to be sensitive to emodepside. Brood size and development were both inhibited in the N2 strain (Kruskal-Wallis, brood size: p = 1.49×10^−42^, animal length: p = 4.32×10^−37^, optical density: p = 1.25×10^−39^), suggesting that the N2 strain is indeed sensitive to emodepside in the HTA. Although development was not affected by emodepside for either mutant strain (Kruskal-Wallis, BZ142 animal length: p = 0.27 and optical density: p = 0.15, and NM1968 animal length: p = 0.60 and optical density: p = 0.12, **Figure 2, Supplementary File 1**), the mutant strains both had higher brood sizes than the N2 strain in emodepside (Kruskal-Wallis, BZ142: p = 4.87×10^−11^ and NM1968: p = 6.97×10^−3^). Brood sizes of these putative null mutant strains were not affected even at high concentrations of emodepside (Kruskal-Wallis, brood size in increasing concentrations of emodepside: BZ142, p = 0.30 and NM1968, p = 0.26). This result confirms that the mutant strains are indeed resistant to emodepside. By contrast, both BZ142 and NM1968 had lower brood sizes than the N2 strain in control conditions, indicating that *slo-1* plays a role in reproduction (**Supplementary Figure 2, Supplementary File 1**). Both deletion strains also had smaller average animal lengths and lower optical densities than the N2 strain in control DMSO conditions (**Supplementary Figure 2, Supplementary File 1**), which again demonstrates that the putative *slo-1* null mutants are less fit than the N2 strain in control conditions. These results recapitulate previous studies and illustrate the applicability of the HTA to study emodepside responses in *C. elegans*.

**Figure 2.**
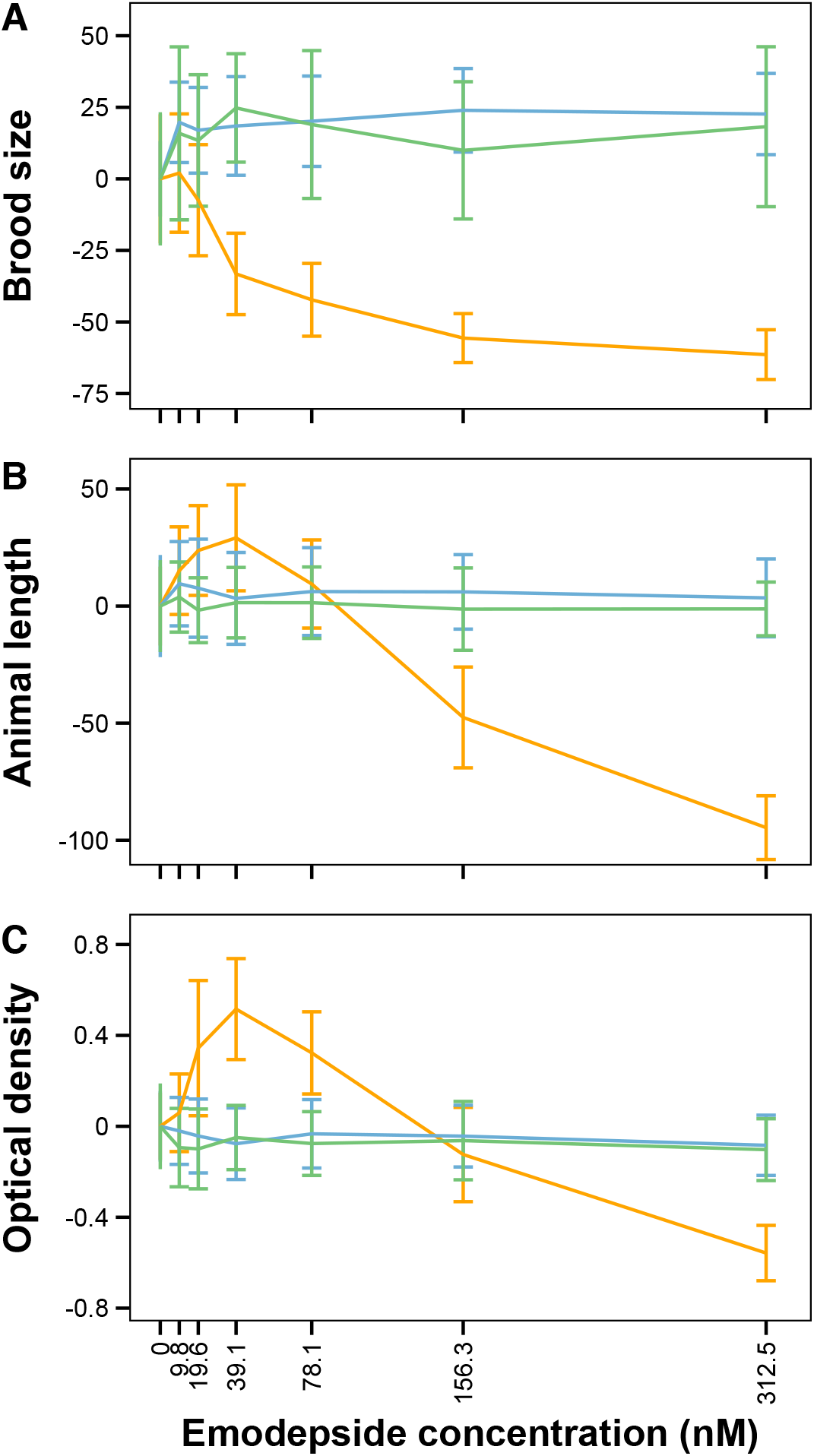
Dose response curves for (A) brood size, (B) animal length, and (C) optical density of the N2 strain (orange) and two *slo-1* putative null mutant strains (BZ142 = blue, NM1968 = green). Phenotypic response values on the y-axis are corrected for the average strain response in control DMSO conditions.

### 3.2 Natural differences in emodepside response are heritable

Previous studies of *C. elegans* resistance to emodepside have been conducted using the N2 strain or mutant strains in the N2 genetic background. Assaying natural variation in *C. elegans* was previously shown to be a powerful tool to identify genetic variation that correlates with differences in benzimidazole responses (Hahnel et al., 2018). To test if the response to emodepside varies by genetic background, we exposed a panel of three genetically divergent *C. elegans* wild strains (N2, CB4856, and DL238) to increasing concentrations of emodepside. At 78.1 nM emodepside, the phenotypic variation was maximized among strains and minimized within replicates of the same strain as shown by broad-sense heritabilities of 88% for brood size, 61% for animal length, and 60% for optical density (**Supplementary Figure 3, Supplementary File 2**). At this concentration, N2 animals were both shorter and less optically dense in the presence of emodepside compared to animals grown in control (DMSO) conditions, showing that development was delayed (**Figure 3A, Supplementary File 3**). The CB4856 and DL238 strains were less affected by this emodepside concentration (**Figure 3B, Supplementary File 3**). These differential responses across the strains and the high heritabilities suggest that genetic factors underlie natural variation in emodepside responses.

**Figure 3.**
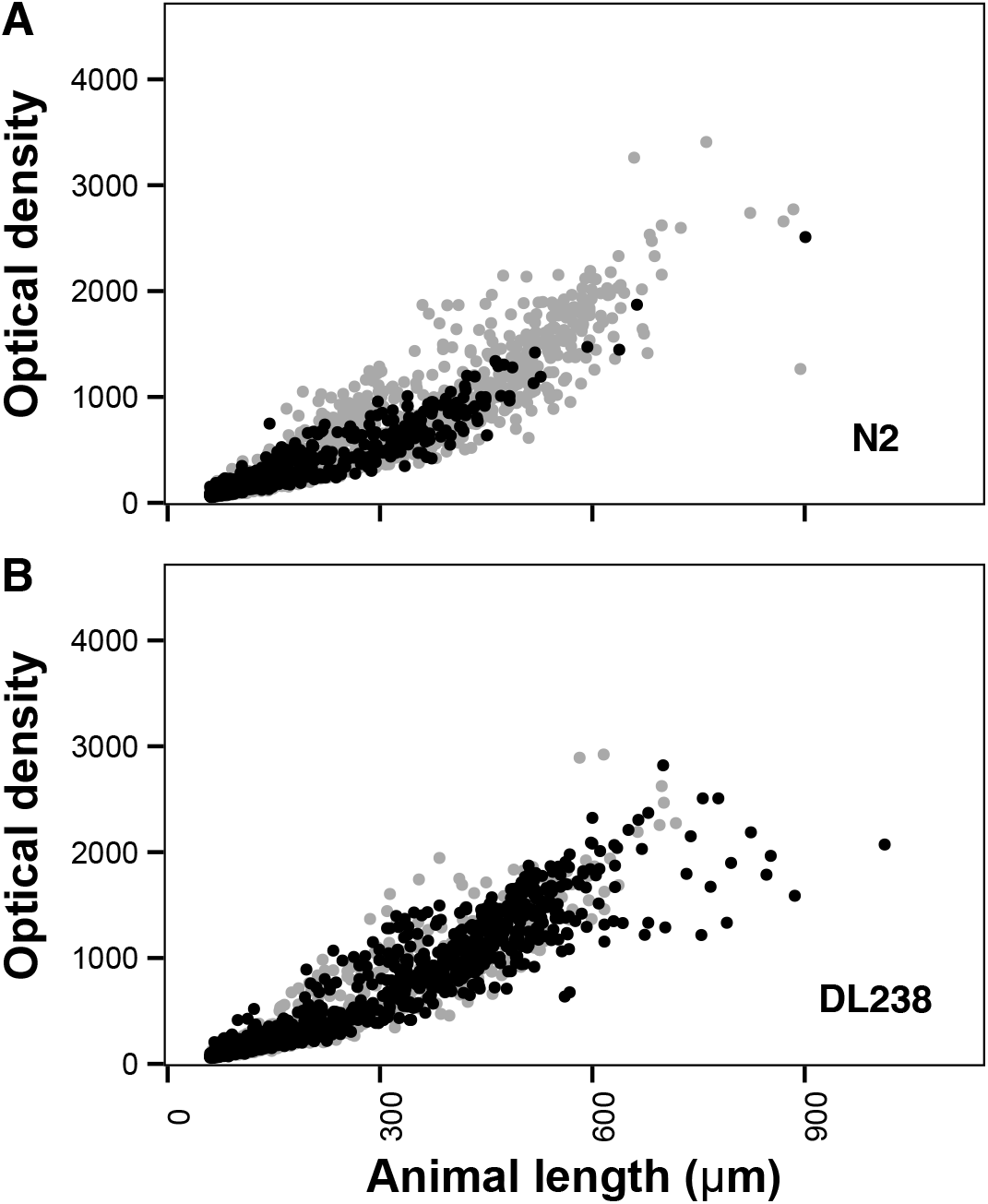
Plots of length and optical density values are shown for each nematode from (A) the sensitive laboratory-adapted strain N2 strain and (B) a more resistant wild strain from Hawaii, DL238. The gray points are nematodes grown in the presence of DMSO (control conditions), and the black points are nematodes grown in the presence of DMSO and 78.1 nM emodepside (emodepside conditions). *C. elegans* grows longer and more optically dense as it ages, and anthelmintic effects can be measured as changes in the demography of animals such that developmental delay is observed as smaller and less dense animals. Here, the DL238 strain was resistant to emodepside because it grew equally well in both control and emodepside conditions.

### 3.3 Emodepside affects brood size and development in a dose-dependent manner

To describe the effects of genetic background on development and brood size in response to emodepside in more detail, we selected nine genetically diverse wild strains and the N2 strain for a second dose response assay to more highly replicate natural differences across the species. We detected significant variation in the dose-dependent responses to emodepside among strains (broad-sense heritabilities at 78.1 nM: 59.5% for brood size, 70.8% for animal length, and 70.8% for optical density). The sensitive laboratory strain N2 falls in the middle of this range, demonstrating that some wild strains are more susceptible to emodepside than the N2 strain and other strains are more resistant (**Figure 4, Supplementary File 4**). At the highest concentration of 312.5 nM emodepside, development and brood size were reduced for all strains. Although high concentrations of emodepside were shown to have detrimental effects on brood size and development, low concentrations of emodepside actually produced larger brood sizes compared to control conditions (**Figure 5, Supplementary Figure 4, Supplementary File 4**). This positive effect on fitness at low concentrations of the drug is called a hormetic effect (Bukowski and Lewis, 2000). For developmental traits the identification of a potential hormetic response is confounded by increased reproduction, because strains that develop further in low concentrations of emodepside start producing a second generation. This second generation increases the observed brood size, but also decreases the average length and optical density of the population because the next generation of early larval stage animals are short and not optically dense (**Supplementary Figure 1**). Regardless of the presence of a hormetic effect for development, the increased brood size at low doses of emodepside suggests emodepside causes a hormetic effect on reproduction.

**Figure 4.**
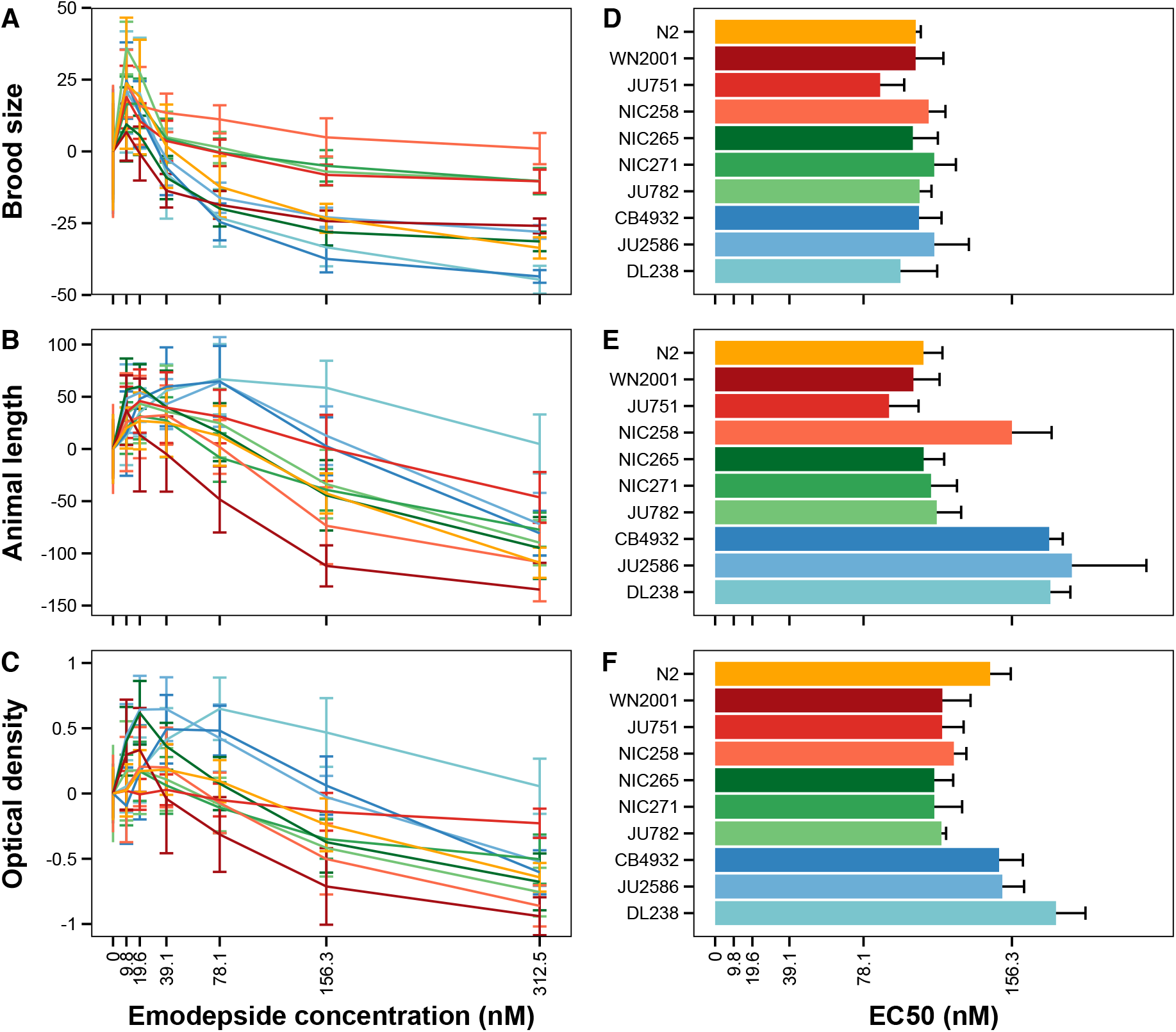
Dose response curves of nine wild *C. elegans* strains and N2 of (A) brood size, (B) animal length, and (C) optical density are shown on the left. Phenotypic response values on the y-axis are corrected for the average strain response in control DMSO conditions. Average EC_50_ values per strain for (D) brood size, (E) animal length, and (F) optical density are shown on the right.

**Figure 5.**
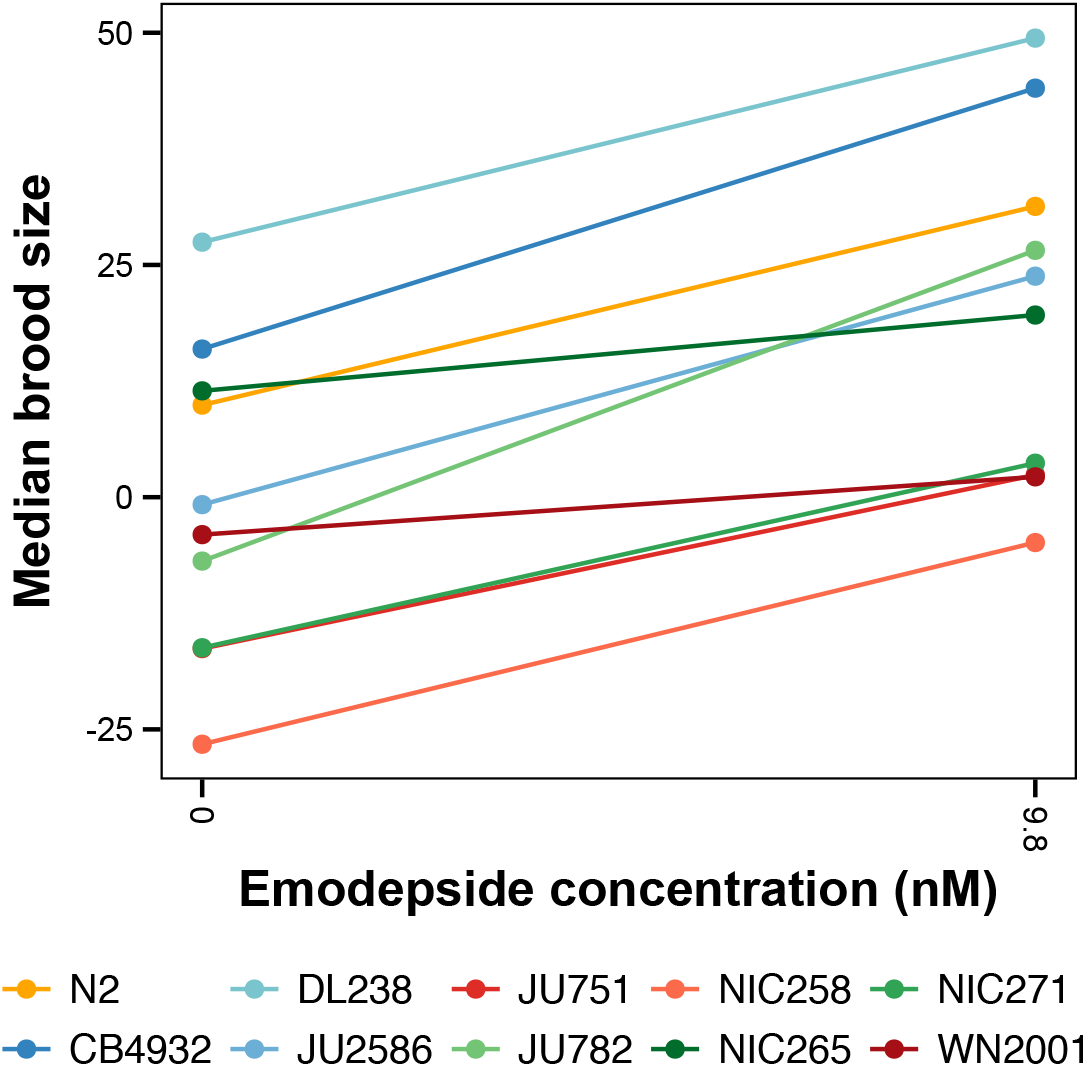
Plot of median brood sizes at the control condition (DMSO) and at 9.8 nM emodepside for ten *C. elegans* strains. Statistical significance of the phenotypic response in DMSO compared to 9.8 nM emodepside was calculated using a pairwise Wilcoxon test. All strains, except NIC265, showed a significant (p < 0.05) hormetic effect.

We next calculated the concentration with half of the maximal drug effect (EC_50_) for each of the strains and all three traits (**Figure 4 D-F, Supplementary File 4**). For all traits, the EC_50_ was significantly affected by genetic diversity across the strains that were assayed (Kruskal-Wallis, brood size: p = 0.0422, animal length: p = 4.56×10^−7^, optical density: p = 1.21^−6^). Overall, these results demonstrate that natural variation in *C. elegans* affects the emodepside response, indicating that this model provides an excellent system to study the genetics of emodepside mode of action and resistance.

### 3.4 Natural variation in candidate gene *slo-1* correlates with resistance in reproduction but not development in wild C. elegans strains

Emodepside has been shown to directly interact with and open the *C. elegans* SLO-1 channel (Kulke et al., 2014), and putative *slo-1* null mutants are resistant to emodepside treatment (**Figure 2**, **Supplementary File 1**(Guest et al., 2007)). Because of these results, we expected that genetic variation in *slo-1* with a predicted moderate or high deleterious effect on gene function would correlate with emodepside resistance. Of nine wild strains assayed here, four (NIC258, NIC265, NIC271, and JU782) have the same four predicted variations in SLO-1 (Arg134Trp, Leu327Phe, Cys328Leu, and Arg678Leu) that causes deleterious amino acid substitutions with a summed BLOSUM score of −6 (Henikoff and Henikoff, 1992). This variation correlated with higher EC_50_ values for brood size but lower EC_50_ values for development (Kruskal-Wallis, brood size: p = 0.0221, animal length: p = 0.263, optical density: p = 3.35×10^−5^).

We also investigated natural variation in another candidate gene for emodepside resistance, *lat-1* (Saeger et al., 2001; Willson et al., 2004). All nine wild strains in the dose response assay harbor natural variants in *lat-1*. To investigate if that variation is predicted to affect *lat-1* function, we summed BLOSUM scores for each of the wild strains. Only the DL238 strain had a negative BLOSUM score (−1), and this score was not correlated with resistance across all strains (Kruskal-Wallis, brood size: p = 0.223, animal length: p = 0.223, and optical density: p = 0.117). In this set of ten strains, variation in *lat-1* does not underlie differences in emodepside responses. Because strains vary in emodepside responses, our results indicate that amino acid variation in *slo-1* or *lat-1* does not explain all differences in emodepside responses, suggesting that additional genetic mechanisms affect the response to emodepside.

## 4. Discussion

Emodepside is a broad range anthelmintic with a distinct mode of action compared to other anthelmintics (Epe and Kaminsky, 2013). Previous studies of emodepside sensitivity and phenotypic effects in *C. elegans* have focussed on the laboratory strain N2 (Bull et al., 2007; Guest et al., 2007; Willson et al., 2004). In this strain, emodepside inhibits egg-laying, pharyngeal pumping, development, and locomotion. In the present study, sensitivity to emodepside was measured across wild *C. elegans* strains, the N2 strain, and two putative *slo-1* null mutants (BZ142 and NM1968) using a large-particle flow cytometer high-throughput assay (HTA) (**Figure 1**). Resistance to emodepside caused by the putative loss-of-function *slo-1* mutants was confirmed using the HTA. Additionally, the effects of emodepside on brood size and development varied across the wild strains (**Figures 3 and 4, Supplementary Files 3 and 4**) and was correlated with protein-coding variation in the resistance candidate gene *slo-1*. Importantly, we found that low doses of emodepside have a hormetic effect on brood size in *C. elegans*. Hormesis was observed in the N2 laboratory strain and eight out of nine wild strains, regardless of their susceptibility to emodepside (**Figure 4, Supplementary File 4**). This consistent hormetic effect across the wild *C. elegans* strains included in our study suggests that emodepside might also cause a hormetic effect in parasitic nematodes. This study shows the power of using natural variation in *C. elegans* to study emodepside responses.

### 4.1 High-throughput assays across wild strains show the effects of emodepside on development and reproduction

In the present study, development and reproductive success in the presence of emodepside was measured for the N2 strain and a set of wild *C. elegans* strains using a HTA. Previously, reproduction was measured on agar plates (Bull et al., 2007), where both the timing and the quantity of egg laying was measured. Using the HTA, brood size is assayed as the overall reproductive success of three L4 larvae over a 96-hour period. Results from agar-based assays and HTA are highly correlated, and HTA intra- and inter-assay correlations are substantially greater compared to agar plate-based methods (Andersen et al., 2015). On agar plates, emodepside prevents egg laying, and the animals are bloated with embryos at higher drug concentrations (20 nM - 500 nM) (Bull et al., 2007). In the HTA, reproduction was inhibited for the N2 strain as well as the wild strains at concentrations similar to previous studies (19.6 μM to 312.5 nM), confirming that brood size in response to emodepside can be measured reproducibly with both assays. Agar-based developmental rate, based on the percentage of eggs that hatch and reach different larval stages in increasing concentrations of emodepside, is delayed (Bull et al., 2007). The HTA measures animal length and optical density of a population established by three L4 larvae over a 96-hour period. The two lowest concentrations of emodepside, 9.8 nM and 19.8 nM, have either no effect on development or a hormetic effect. Higher concentrations (39.1 μM to 312.5 nM), which overlap with the effective concentrations from the agar-based development phenotypes, negatively affect animal length and optical density (**Figure 4**). The results from both the agar-based and HTA methods indicate that emodepside inhibits reproduction at lower concentrations than development. Emodepside inhibited reproduction from approximately 20 nM and up, compared to approximately 40 nM and up for development. The agar-based study did not find a hormetic effect, but our results suggest that such an effect is likely to be present at concentrations below the range tested on agar plates. Our results show that the HTA provides a platform to screen hundreds of strains efficiently and that the different measures of reproduction and development are similarly affected across assay platforms.

### 4.2 Natural variation affects development and brood size in the presence of emodepside

The response of *C. elegans* to emodepside is affected by natural genetic variation (**Figure 4, Supplementary File 4**). Our results showed that all strains are affected by emodepside, and that higher doses inhibit development, as measured by animal length and optical density, and reproduction, as measured by brood size (**Figure 4, Supplementary File 4**). For brood size, strain-specific differences were correlated with variation in *slo-1* where strains with predicted deleterious variation were more resistant to emodepside treatments. However, strains with higher brood sizes at lower concentrations do not have variation in *slo-1*, suggesting that the hormetic effect is not mediated by *slo-1*. It will be informative to introduce *slo-1* variation in the wild strains with higher fitness using CRISPR-Cas9 genome editing to test if *slo-1* variation reduces brood size in a more resistant background. Our results show that reproduction and development are inhibited by higher concentrations of emodepside, and that natural variation affects the extent of this response. Future measurements of emodepside responses across additional wild isolates will improve the power to detect candidate resistance genes across the species using genome-wide association studies.

### 4.3 Hormetic effects of emodepside suggest a potential risk of treatment failure at suboptimal doses

Nine of ten strains showed a hormetic response in reproduction when treated with a low emodepside concentration (**Figure 4 A-C, Figure 5, Supplementary File 4**). The presence of hormetic responses across strains illustrates that hormesis is a common response to low concentrations of emodepside across *C. elegans* strains. If this response is shared with parasitic nematodes, then experimental designs to calculate EC_50_ values in parasites need to account for this effect. A low EC_50_ estimate caused by a hormetic effect can lead to recommended treatment doses that will be insufficient to treat parasitic nematode infections. Underdosing is a known risk factor for the selection of resistance against all anthelmintics (Sangster et al., 2018; Silvestre et al., 2001; Smith et al., 1999). If hormetic doses are administered, either because of erroneous EC_50_ calculations or as a result of other treatment factors like ineffective drug delivery (Sangster et al., 2018), the infection might be intensified rather than treated. Our results also imply that low doses of emodepside are beneficial rather than detrimental for nematode growth. To prevent hormesis from rendering emodepside treatment ineffective, it is essential to investigate hormetic effects and adjust treatment recommendations accordingly.

### 4.4 Natural variation in *C. elegans* can facilitate the study of anthelmintic resistance

The free-living nematode *C. elegans* is a long standing model to study anthelmintic mode of action and resistance of parasitic nematodes (Geary and Thompson, 2001; Hahnel et al., 2020; Holden-Dye et al., 2014; Wit et al., 2020). The suitability of *C. elegans* as a model is the result of a range of attributes, including the phylogenetic relationship of *C. elegans* with many parasitic nematodes of human and veterinary importance, its short and direct life cycle, a wide range of genome-editing tools, and its high-quality reference genome and gene models. Additionally, larval stages of many parasitic nematodes occupy the same niches as *C. elegans* (Crombie et al., 2019; Frézal and Félix, 2015). Similar environmental stressors, including naturally occuring precursors of anthelmintics (Alivisatos et al., 1962; Campbell, 2012, 2005), can cause similar selective pressures for both species to evolve resistance.

Previous studies on the emodepside resistance candidate gene *lat-1*, showed that putative *lat-1* null mutants were resistant in reproduction and pharyngeal pumping assays, but sensitive in locomotion assays (Guest et al., 2007), and that putative *slo-1* null mutants are resistant in all assays. Here, we show that although putative *slo-1* null mutants are resistant to emodepside treatment, *slo-1* variation is not the only determinant of resistance across wild strains. These results imply that multiple genes likely affect the response to emodepside. To identify these genes, genetic variation across wild strains can be correlated with phenotypic responses to emodepside. Genes identified based on population-wide variation are more likely to translate to other species than genes identified based on one genetic background. After identification of candidate genes, genetic variation in these genes should be tested in a controlled genetic background by introducing specific mutations using CRISPR-Cas9 genome editing (Dilks et al., 2020).

## Footnotes

1 Note: Supplementary data associated with this article.

## Acknowledgements

We would like to thank members of the Andersen laboratory for their technical assistance and helpful comments on the manuscript. This work was supported by R01 AI153088 to E.C.A. We would also like to thank Wormbase and the *C. elegans* Natural Diversity Resource (CeNDR) for data and tools critical for our analyses of natural variation (NSF CSBR 1930382). The *slo-1* mutant strains (BZ142 and NM1968) were provided by the *Caenorhabditis* Genetics Center, which is funded by NIH Office of Research Infrastructure Programs (P40 OD010440).

