## Supplementary Figures for "Natural variation in *Caenorhabditis elegans* responses to the anthelmintic emodepside"

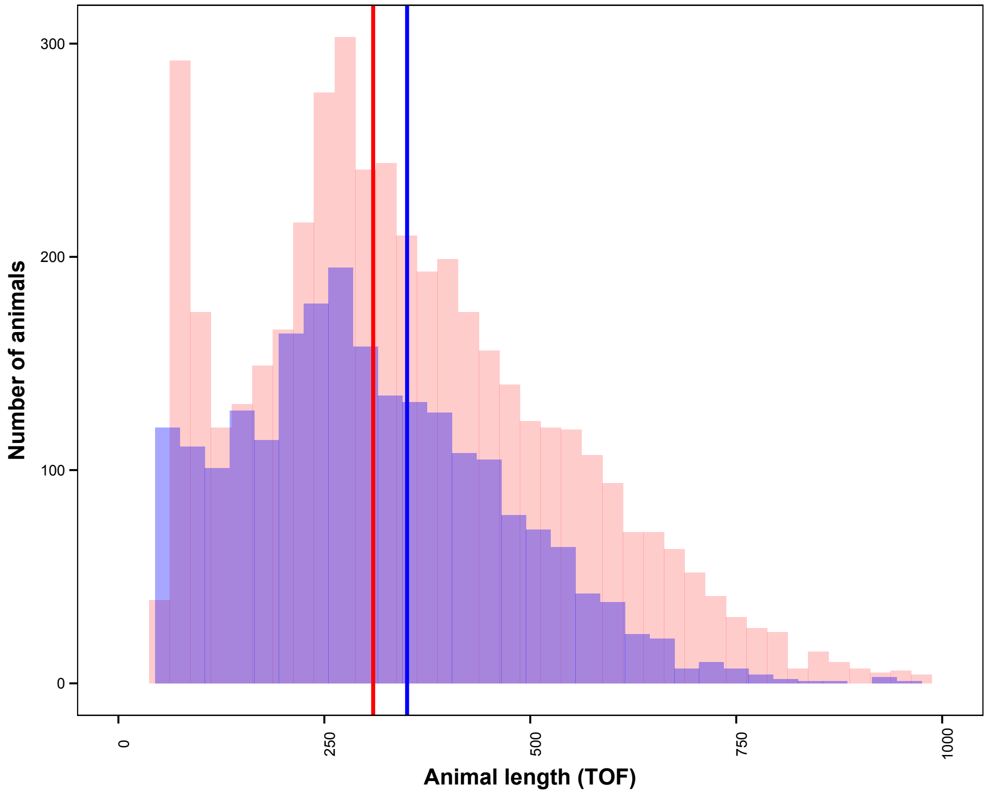


**Supplementary Figure 1:** Emodepside resistance is determined as a function of nematode brood size and development using the previously developed high-throughput assay (HTA). In this assay, three L4 larvae grow to adults and produce offspring for 96 hours. These offspring are laid and hatch at different times during this 96-hour assay, so they range in size from small early larval stages, which hatched late in the assay, to large late larval stages, which hatched early in the assay. Near the end of 96 hours, some offspring that hatched early in the assay will develop into adults and lay embryos that hatch into early small larvae. After 96 hours, the total number of progeny is measured by counting all objects in a well. This number includes progeny from the original three parents but can sometimes include the additional progeny from the next generation. An emodepside sensitive strain (blue) will have fewer offspring than a resistant strain (red). The brood size can be overestimated for resistant strains because the offspring that hatch late in the 96-hour assay are also counted. After 96 hours, the lengths of all objects in a well are measured.

The length measurements for the entire population can be shown as a distribution. Progeny laid early in the 96-hour assay will be longer than progeny laid late in the assay. We use summary statistics (*e.g.* mean) to describe the length distribution. Many times, an emodepside sensitive strain (blue) will have a mean length larger (vertical blue line) than a resistant strain (vertical red line) because the offspring that hatch late in the 96-hour assay are small and push the average down.

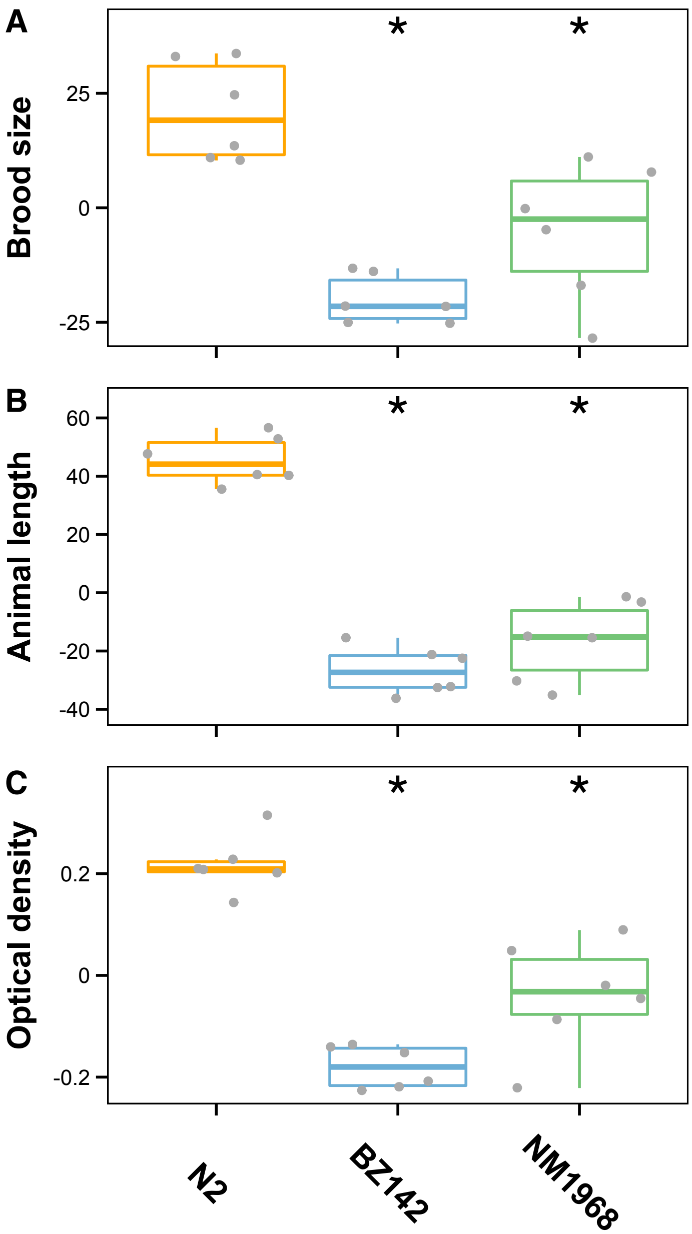


**Supplementary Figure 2:** (A) Brood size, (B) Animal length, and (C) Optical density of the N2, BZ142, and NM1968 strains in control (DMSO) conditions (y-axis) are plotted as Tukey box plots by strain (x-axis). The horizontal line in the middle of the box is the median, and the box denotes the 25th to 75th quantiles of the data. The vertical line represents the 1.5x interquartile range. Statistical significances of BZ241 and NM1968 compared to the N2 strain calculated using a pairwise Wilcoxon test are shown above each strain, * = p-values < 0.05 with Bonferroni correction.

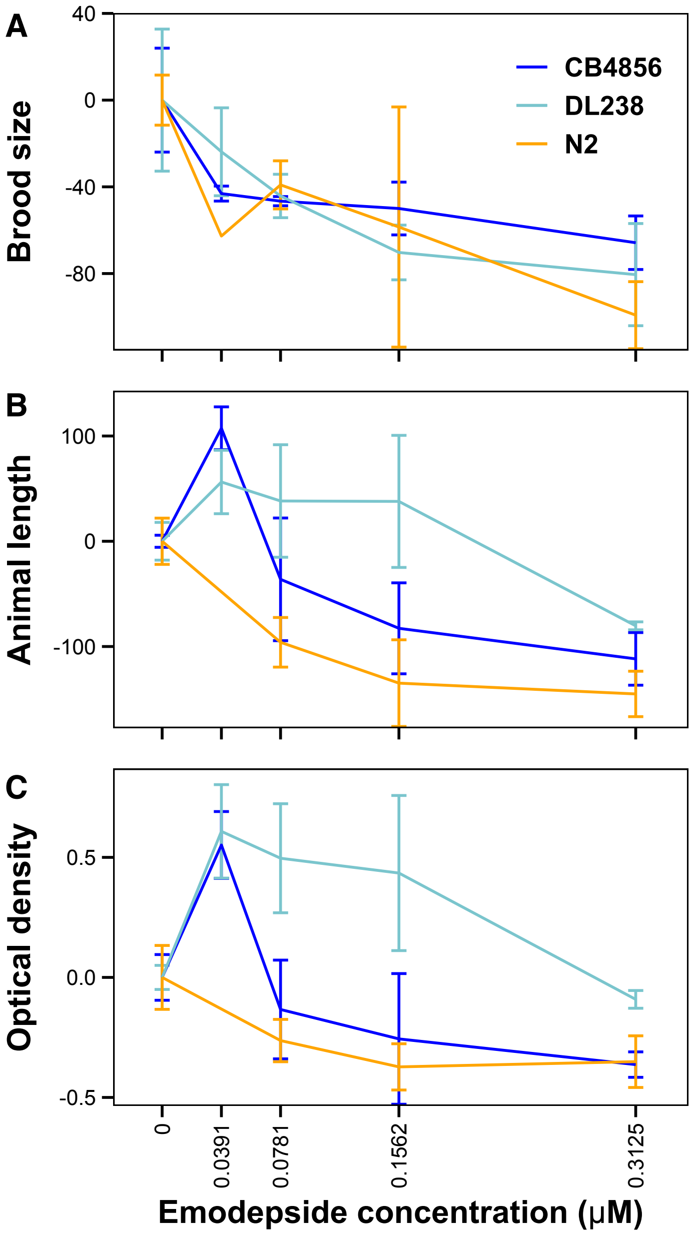


**Supplementary Figure 3:** Dose response curves for (A) brood size, (B) animal length, and (C) optical density.

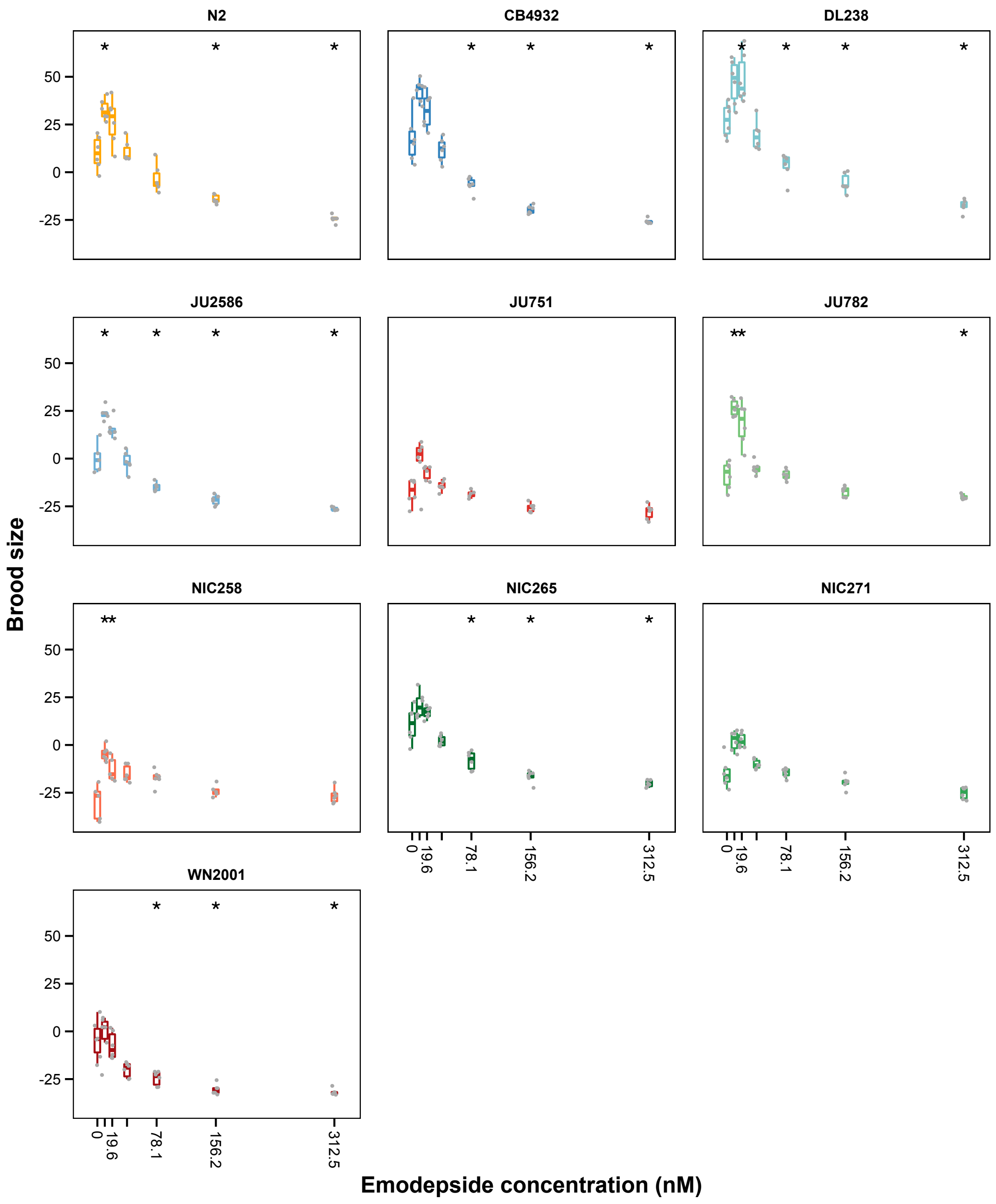


**Supplementary Figure 4**

Brood sizes (y-axis) are plotted as Tukey box plots by emodepside concentration (x-axis). The horizontal line in the middle of the box is the median across six technical replicates, and the box denotes the 25th to 75th quantiles of the data. The vertical line represents the 1.5x interquartile range. Statistical significance of increasing emodepside concentrations is calculated with a pairwise Wilcoxon test with DMSO as reference group and shown above the boxes, * = p-values < 0.05 with Bonferroni correction.
